# DSSP 4: FAIR annotation of protein secondary structure

**DOI:** 10.1101/2025.04.11.648460

**Authors:** Maarten L. Hekkelman, Daniel Álvarez Salmoral, Anastassis Perrakis, Robbie P. Joosten

## Abstract

Protein secondary structure annotation is essential for understanding protein architecture, serving as a cornerstone for structural classification, alignment, visualisation, and machine learning applications. The Define Secondary Structure of Proteins (DSSP) algorithm has long been the standard for assigning secondary structure elements such as α-helices, β-sheets, and loops in protein models. Here, we introduce DSSP version 4, which recapitulates DSSP functionality in a modern computational framework, extending also to the detection of left-handed κ-helices (Poly-Proline II helices). To align with the FAIR principles (Findable, Accessible, Interoperable, Reusable), DSSP 4 adopts mmCIF as its primary input and output format, while retaining compatibility with legacy PDB and DSSP formats. We applied this updated tool to analyse the distribution of secondary structure elements across the Protein Data Bank (PDB) differentiating structures from diverse experimental methods, revealing insights into the prevalence and length of secondary structure elements, including the newly annotated κ-helices. The DSSP 4 software, databank, and server are freely accessible from https://pdb-redo.eu/dssp, ensuring broad utility and interoperability in structural biology research.

## 1. Introduction

Protein secondary structure plays a vital role in describing and illustrating the structure of proteins, being a cornerstone of protein visualisation in molecular graphics. Moreover, secondary structure is used to classify and cluster protein structure families (Murzin et al. 1995; Orengo et al. 1997) and as a basis for protein structure alignment (Krissinel and Henrick 2004). The annotation of secondary structure elements, such as α-helices, β-sheets, and loops is also a key step in the analysis of structure models or ensembles (Lazar et al. 2021; Herrera et al. 2025). It can also be used to compare different structure modelling procedures, e.g. those used to predict the outcome of mutations (Pan et al. 2022). Per-residue assignment of secondary structure nowadays is also an important feature for diverse machine learning purposes. It was for instance used to train the ESM3 language model that was e.g. applied to design new fluorescent proteins (Hayes et al. 2025) and to train a hierarchical graph neural network to predict nucleic acid binding residues in proteins (Xia et al. 2021).

Most structural biologists can confidently identify α-helices and β-strands in protein structure models without the aid of dedicated annotation tools. However, when less common secondary structure elements such as 3_10_- and π-helices are involved or when the elements must be delineated exactly, manual identification becomes error-prone. The problem of secondary structure annotation is indeed “trivial, but difficult” (Richardson and Richardson 1989) and to achieve consistency, particularly over large sets of structure models like the Protein Data Bank (PDB) (wwPDB consortium 2019) and the AlphaFold Protein Structure Database (AFDB) (Varadi et al. 2022), an algorithmic approach is needed. Over the years, many such algorithms were introduced (Frishman and Argos 1995; Labesse et al. 1997; King and Johnson 1999; Taylor 2001; Andersen et al. 2002; Majumdar et al. 2005; Martin et al. 2005; Law et al. 2014), each with their own approach and resulting strengths and weaknesses. The wealth of algorithms also lead to servers like 2Struc that allows running a multitude of tools and combine their different annotations (Klose et al. 2010). However, the Define Secondary Structure of Proteins (DSSP) algorithm (Kabsch and Sander 1983) was an early algorithm to assign secondary structure and has grown to become the *de facto* standard.

The DSSP software and the databank of DSSP-annotated PDB entries have been kept available over decades (Joosten et al. 2011; Touw et al. 2015) with several full software re-implementations to keep up with the changing landscape in computational structural biology while keeping the annotation algorithm consistent. Only minor changes were made to detect previously overlooked π-helical turns in α-helices (Cooley et al. 2010) (also referred to as α-bulges (Van der Kant and Vriend 2014) or π-bulges (Cartailler and Luecke 2004)). Here we describe the newest DSSP version 4 with new functionality to detect left-handed Poly-Proline II (PPII) helices which are commonly occurring conserved structural elements (Adzhubei and Sternberg 1993; Adzhubei and Sternberg 1994; Cubellis et al. 2005; Adzhubei et al. 2013). Due to the fact that they need not contain any Proline residues, we will refer to them as κ-helices (Meirson et al. 2020). In addition, the DSSP software and databank were upgraded to now use mmCIF (Westbrook et al. 2022) as the primary input and output format to deal with the requirement of FAIR (Findable, Accessible, Interoperable and Reusable) resources in structural biology, while keeping support for the legacy PDB and DSSP formats for input and output, respectively.

Using the new DSSP software we analysed the distribution of secondary structure elements in the Protein Databank, the AlphaFill databank that contains predicted structure models, and the metagenomic protein models in the NMPFamsDB (Pavlopoulos et al. 2023; Baltoumas et al. 2024). We also analysed the distribution of secondary structure elements in structure models derived from different experimental techniques.

## 2. Results and discussion

### 2.1 Detection of κ-helices (PPII helices)

When the original DSSP was written, there were very few κ-helices in the PDB. Moreover, κ-helices do not have a specific hydrogen bonding pattern that would allow detection by the DSSP algorithm similar to how other secondary structure elements are detected. As a result, detection of κ-helices was not implemented. The PDB is now over a thousand times larger with ample examples to add a detection algorithm. In the current DSSP software, κ-helices are marked when there are three consecutive residues with a φ-torsion angle of -75±29° and a Ψ-torsion angle of 145±29°, based on de criteria of Mansiaux *et al*. (Mansiaux et al. 2011). An additional requirement is that and no other secondary structure type is already assigned. The minimal length of κ-helices can be changed from the command line. The DSSP per-residue alphabet was extended with ‘P’ marking a κ-helix (Table 1). Conveniently, the legacy DSSP format had space to annotate κ-helices without breaking backward compatibility.

**Table 1:** DSSP secondary structure types and their relative occurrence.

| **Table 1: DSSP secondary structure types and their relative occurrence** |  |  |
| --- | --- | --- |
| **DSSP code** | **Description** | **%-age in DSSP databank** |
| B | Residue in isolated β-bridge | 1.2 |
| E | Extended strand, participates in $\beta$ ladder | 21.5 |
| G | $3_{10}$-helix | 3.6 |
| H | α-helix | 32.3 |
| I | π-helix | 0.5 |
| P | κ-helix (PPII helix) | 1.9 |
| S | Bend | 8.9 |
| T | Hydrogen-bonded turn | 11.1 |
| . | No distinct secondary structure; loop | 19.0 |

Using the newly added functionality we updated the DSSP databank. The DSSP databank covers the whole PDB and is updated automatically when new or updated PDB entries are released (see methods). For the analyses we took a snapshot of the DSSP databank on February 7^th^ 2025 (with 1,874,031 entries) and checked the relative frequency of different secondary structure elements (Table 1). Residues in α-helices and β-sheets are the most common with 32.3% and 21.5%, respectively. The newly implemented κ-helices, comprised 1.9% of all residues. As the PDB and by extension the DSSP databank contain many (close) homologs, this number will be biased towards proteins that are common in the PDB. A more representative subset of the PDB (16k chains with less than 25% mutual sequence identity) defined by the PISCES server (Wang and Dunbrack 2003) had 2.0% of the residues as part of a κ-helix, which is lower than the values described in Mansiaux *et al*. (Mansiaux et al. 2011). We also analysed the length of κ-helices in the PDB. Rather than using the one-letter codes which summarises all the independent secondary structure assignments, we used the κ-helix-specific annotation (see the Methods section). This annotation, in which for each residues it is assigned whether it is part of a minimal helical element also exist for the other helix types (Kabsch and Sander 1983). As these annotations are independent, a residue can be assigned to more than one helix type, which happens particularly at the start of helices. κ-helices are typically very short with half of them being three residues or shorter and 90% four residues or shorter. The longest κ-helix, 48 residues, is found in PDB entry 8tx1 which is a filamentous part of a mastigoneme in the flagella of *Chlamydomonas reinhardtii* (Wang et al. 2023). This long helix is followed by three shorter κ-helices (of 4, 14, and 8 residues respectively) that are part to same extended structure. PDB entry 8tx1 is determined by cryo-electron microscopy at 3.6Å resolution. Most other very long helices (of any type) are also found in structure models determined by cryo-EM.

### 2.2 Secondary structure elements for different sources of structure models

The choice of experimental technique used to obtain a structure model of a protein depends on the sample under consideration. For instance, X-ray crystallography works very well for proteins and complexes of varying sizes if they can be crystallised. Cryo-EM does not require crystallisation and works very well for large and very large complexes and complexes containing nucleic acids, but is very challenging (or impossible) for small proteins. NMR is very good at capturing protein dynamics but cannot be used for structure determination of large proteins. Another notable difference is the way models are made: in X-ray crystallography and Cryo-EM it is common practice to not model residues with poor electron density/potential maps, whereas in NMR typically whole proteins are modelled and limitations of the experimental data are reflected in wider ensembles of conformations. Figure 1 shows the distribution of secondary structure elements in the DSSP databank broken down by experimental technique. A notable difference between X-ray crystallography and Cryo-EM is the percentage of α-helix and β-strand residues: X-ray crystallography has a relatively high fraction of β-strand (22% vs. 18%) but a low fraction of α-helix (32% vs. 36%) compared to cryo-EM. The proteins studied with neutron diffraction have a high percentage of 3_10_-helical residues (6.5%) versus the DSSP-wide percentage of 3.6%. NMR and electron diffraction both have more than 50% of their residues in the poorly ordered classes of turns, bends and loops. In NMR this is likely the result of how structure models are made, whereas for electron crystallography this may depend more on the sample type (microcrystals) and data quality (relatively low data completeness). Percentages for all experimental techniques and secondary structure types are given in Supplemental data 1.

**Figure 1:**
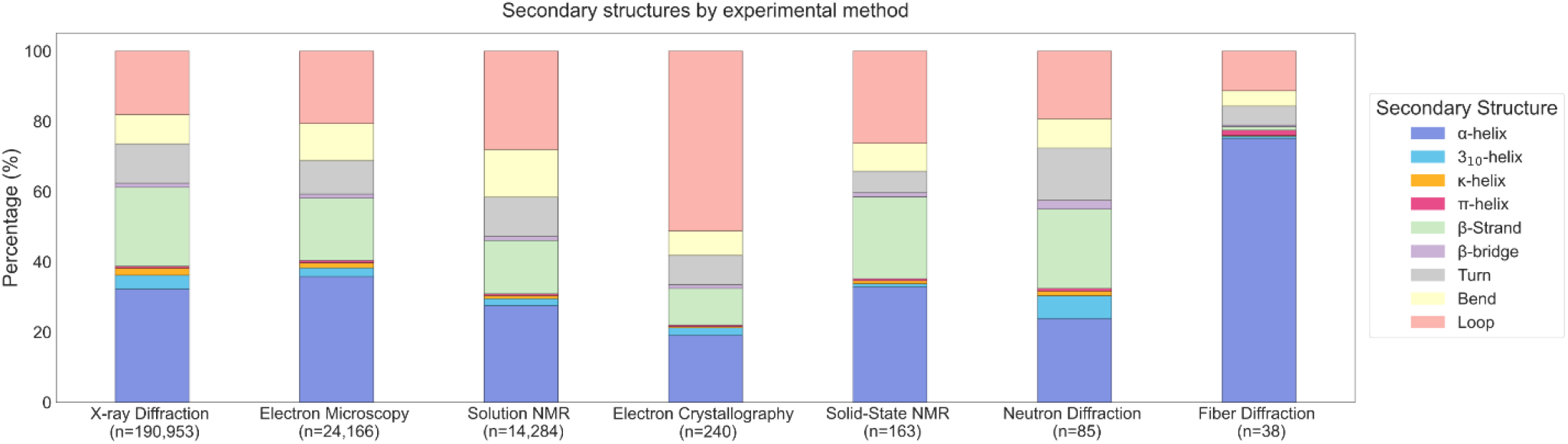
Secondary structure composition in the DSSP databank by experimental method. Stacked bar plots show the percentage of different secondary structure elements grouped by the experimental method used for structure determination.

We also compared experimentally derived structure models to models from AI predictions (Figure 2). For that, we first used predicted models from the AlphaFill databank of ligand supplemented predicted structure models (Hekkelman et al. 2023) which covers predictions from various sources: model organism proteomes; global health proteomes; the SwissProt subset of the AlphaFold Protein Structure Database (AFDB); all entries from the Big Fantastic Virus Database (BFVD) (Kim et al. 2025); and entries that were added on-demand from users. We also used the NMPFamsDB resource of metagenomic structures with novel folds, to assess whether these folds that are unique to metagenomes have a particular distribution of secondary structure elements. Both sources of predicted models have more α-helical content than the PDB (35.6% and 38.7%, compared to 32.3% for the PDB). In terms of β-sheet content, the AlphaFill models only have 15.7%, whereas PDB and NMPFamDB models have 21.5% and 23.6%. Another notable difference is that the amount of loop residues in AlphaFill is much higher (26.1%) than in the PDB (19.0%) and NMPFamsDB (17.0%). This is likely caused by different “filtering” practices. In experimental structures, poorly ordered regions like loops are often not modelled due to poor experimental data. In NMPFamsDB, poor models (pTM score < 0.7) were left out (Pavlopoulos et al. 2023) whereas in the AFDB (and thus in AlphaFill) there is no such filtering.

**Figure 2.**
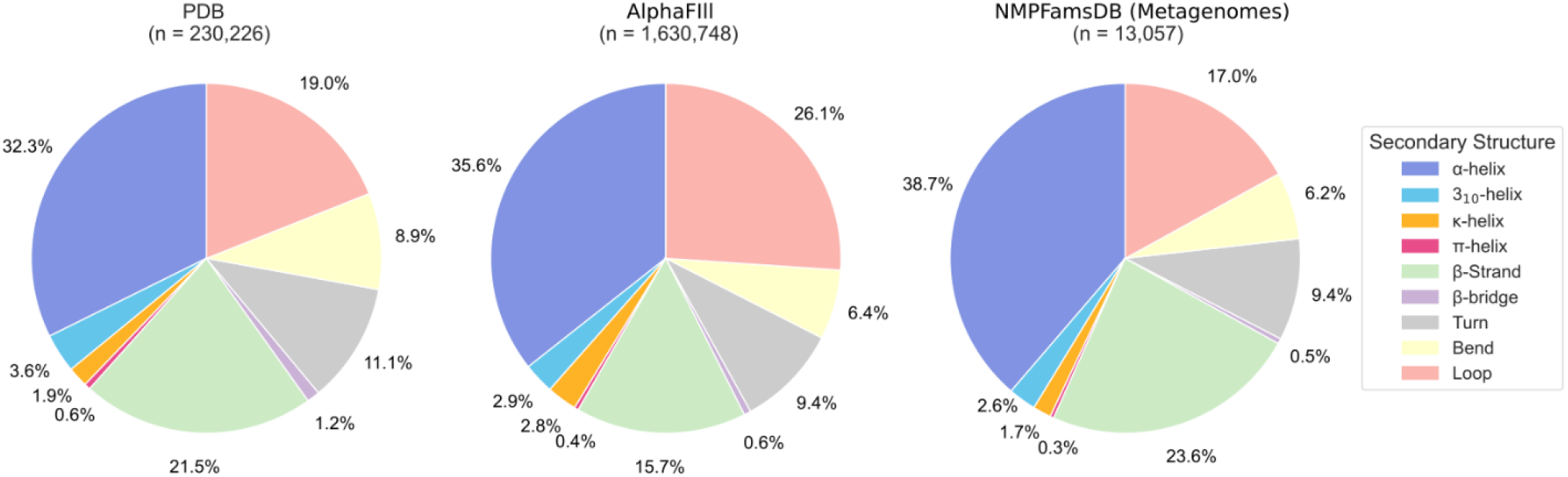
Secondary structure composition: Experimental (PDB) vs. AI-predicted models. Pie charts compare the percentage of secondary structure elements between experimentally determined structures (PDB) and AI-predicted models from AlphaFill and NMPFamsDB (Metagenomes).

### 2.3 Moving on from legacy DSSP format to mmCIF

When the original DSSP software was designed, the column-based PDB format (with very limited standardisation) was the carrier of macromolecular structure data. DSSP’s output followed a similar column based format (Kabsch and Sander 1983) which suited the needs and standard practices of the structural biology community at the time. In FAIR terms, the DSSP format could be considered a community standard and therefore Interoperable and Reusable.

Nowadays, the advances in experimental structural biology, particularly with respect to the size of the macromolecular complexes we can determine, have rendered the PDB format and by extension the DSSP format obsolete. In 1983, no structure models contained more than 1,600 residues whereas there are now cryo-EM entries in the PDB of more than 160,000 residues, e.g. the structure of the sea urchin sperm tail doublet microtubule (PDB entry 8snb) (Leung et al. 2023). Apart from size limitations, the interconnection of structural biology resources also requires data to have a well-documented machine-readable description to allow straightforward Interoperability and Reusability. The mmCIF format, underpinned with a formal dictionary, fits this requirement and is rapidly becoming the standard in structural biology. We therefore made mmCIF the primary input and output format of DSSP. As we also realise that over the years many established workflows exist in the structural biology community that rely on the PDB and DSSP format, DSSP 4 can still read valid PDB files and write the legacy DSSP format. We do however encourage new workflows to use mmCIF and existing workflows to be updated to stay Interoperable and Reusable for the foreseeable future.

All the data from the original DSSP format are written in categories in a DSSP namespace:

*_dssp_struct_summary* for the core DSSP table with per-residue descriptors, *_dssp_struct_bridge_pairs* to describe individual β-bridges, *_dssp_struct_ladder* to describe β-sheets, and *_dssp_statistics*,

*_dssp_statistics_histogram* and *_dssp_statistics_hbond* for summary statistics about e.g. the size of secondary structure elements and about the types of backbone hydrogen bonds. An mmCIF dictionary extension with a formal description of the new records is provided as Supplemental Data 1. The existing

*_struct_conf, _struct_conf_type, _struct_sheet, _struct_sheet_range* categories in the input mmCIF file are reannotated to describe all the secondary structure elements which is particularly useful for display purposes in molecular graphics.

Annotation of a (compressed) structure model of 1,600 residues takes fewer than three seconds on a modest PC. Extremely large structure models like the 160,000 residue microtubule model can take several hours, which is why ready-made DSSP annotations of the PDB are available via the DSSP databank (see below).

### 2.4 Availability

The DSSP resource consists of three elements: the DSSP program called *mkdssp*, the DSSP databank, and the DSSP server that allows users to run DSSP online and access the databank via a web browser. All elements are freely accessible. The code for *mkdssp* is available from https://github.com/PDB-REDO/dssp with a permissive BSD-2-clause licence. The program can also be deployed from the Debian repository (package name *dssp*) or, for a Python environment, through conda-forge. The software is also integrated into the CCP4 software suite (Agirre et al. 2023).

The whole DSSP databank, i.e. annotations of all PDB entries, is available through the *rsync* protocol at rsync://rsync.pdb-redo.eu/dssp/. Individual entries can be downloaded directly from https://pdb-redo.eu/dssp/db/{PDBID}/mmcif(mmCIFformat) and https://pdb-redo.eu/dssp/db/{PDBID}/legacy(DSSPformat). Please note that DSSP formatted files are not available for PDB entries that cannot be cast to the DSSP format, most commonly due to size restrictions. DSSP annotations for AFDB entries can be found directly in the mmCIF versions of the AFDB entries or in the downstream AlphaFill models. Similarly, DSSP annotations of BFVD entries are available through AlphaFill. DSSP annotations of NMPFamsDB entries are available via the NMPFamsDB website.

The DSSP server at https://pdb-redo.eu/dssp runs as a single application that serves the website and orchestrates the updates of the DSSP databank (see methods). On the DSSP website users can download DSSP databank entries in both mmCIF and legacy format, and users can upload their own structure models to be annotated. Non-interactive annotation of user provided models is also available although we recommend running the *mkdssp* application locally for annotating large sets of structure models. The code for the DSSP server application is available from https://github.com/PDB-REDO/dssp-server with a BSD-2-clause license.

## 3. Conclusion

The new DSSP reads and writes mmCIF to provide FAIR secondary structure annotations of structure models. For experimental structure models, the secondary structure content differs between experimental methods reflecting both the underlying samples and standard model making practices of the method. Secondary structure content of AI generated models depends strongly on the choice of filtering of the AI output.

## 4. Methods

### 4.1 Creation of the new DSSP software and databank

The DSSP code was rewritten in the C++20 standard using the open source *libcifpp* library(Hekkelman 2025) to provide mmCIF support. Detection of κ-helices was added based on de criteria of Mansiaux *et al*.(Mansiaux et al. 2011) New mmCIF categories were designed to capture DSSP annotations in mmCIF and were formalised in a dictionary extension (Supplemental Data 1). A new DSSP server application was written in C++20. The new application serves the DSSP website and lets users run the DSSP algorithm on uploaded structure models. The server application also replaces the previous make-based maintenance of the DSSP databank(Joosten et al. 2011). Using a local copy of the Protein Data Bank, the server application checks for new or updated PDB entries and stages them for incorporation into the DSSP databank. Following the PDB’s update cycle, the DSSP databank is updated on a weekly basis.

### 4.2 Analysis of secondary structure in different structural resources

Snapshots were taken of the DSSP, AlphaFill and NMPFamsDB resources on February 7^th^ 2025. A predefined subset of the PDB (X-ray and EM entries, better than 4Å resolution, with less than 25% mutual sequence identity) was taken from the PISCES server. This set only listed protein chains determined by X-ray crystallography or Cryo-EM at resolution of 4Å or better that had less than 25% mutual sequence identity. For consistency, all models in PDB-REDO, AlphaFill and NMPFamsDB were analysed with the same DSSP version as is used for the DSSP databank. DSSP annotations of all models were analysed with a custom Python script using the Pandas (The pandas development team 2020) and Numpy packages (Harris et al. 2020). For the analysis of the distribution of κ-helix lengths, the mmCIF item *helix_pp* was taken from the *_dssp_struct_summary* category and used to find κ-helices. Briefly, a helix starts at the first ‘>’ value of a stretch of residues and ends at the first ‘<’ that is followed by an unassigned residue (value ‘.’), by the start of a new helix, or at the end of a chain.

For calculating the percentages of different secondary structure types the *_dssp_struct_summary*.*secondary_structure* item, i.e. the well-known DSSP one-letter code, was used. In the DSSP databank entries the *_exptl*.*method* item was used to determine the experimental source of the structure models.

The figures were generated using the Python modules Matplotlib (Hunter 2007) and Seaborn (Waskom 2021).

## Supplementary materials

supplemental_data1_dssp-extension.dic: An extension to the mmCIF-PDBx dictionary that describes all DSSP categories and items in the mmCIF-formatted DSSP output.

supplemental_data2.csv: Percentages of secondary structure elements per experimental technique or computational data source.

## Author contributions

MLH wrote and maintains the current DSSP code, databank and webserver. DAS performed the data analysis and made figures, AP helped conceptualise and write the manuscript, RPJ conceptualised and wrote the manuscript, and supervises the DSSP development and maintenance.

## Acknowledgements

The authors thank the Research High-Performance Computing facility of the Netherlands Cancer Institute for maintaining DSSP’s compute infrastructure. This work was supported by an institutional grant of the Dutch Cancer Society and of the Dutch Ministry of Health, Welfare and Sport. We also thank CCP4 and Oncode Accelerator for providing financial support.

## References

Adzhubei AA, Sternberg MJE. 1993. Left-handed Polyproline II Helices Commonly Occur in Globular Proteins. Journal of Molecular Biology. 229(2):472–493. doi:10.1006/jmbi.1993.1047.

Adzhubei AA, Sternberg MJE. 1994. Conservation of polyproline II helices in homologous proteins: Implications for structure prediction by model building. Protein Science. 3(12):2395–2410. doi:10.1002/pro.5560031223.

Adzhubei AA, Sternberg MJE, Makarov AA. 2013. Polyproline-II Helix in Proteins: Structure and Function. Journal of Molecular Biology. 425(12):2100–2132. doi:10.1016/j.jmb.2013.03.018.

Agirre J, Atanasova M, Bagdonas H, Ballard CB, Baslé A, Beilsten-Edmands J, Borges RJ, Brown DG, Burgos-Mármol JJ, Berrisford JM, et al. 2023. The CCP4 suite: integrative software for macromolecular crystallography. Acta Cryst D. 79(6):449–461. doi:10.1107/S2059798323003595.

Andersen CAF, Palmer AG, Brunak S, Rost B. 2002. Continuum Secondary Structure Captures Protein Flexibility. Structure. 10(2):175–184. doi:10.1016/S0969-2126(02)00700-1.

Baltoumas FA, Karatzas E, Liu S, Ovchinnikov S, Sofianatos Y, Chen I-M, Kyrpides NC, Pavlopoulos GA. 2024. NMPFamsDB: a database of novel protein families from microbial metagenomes and metatranscriptomes. Nucleic Acids Research. 52(D1):D502–D512. doi:10.1093/nar/gkad800.

Cartailler J-P, Luecke H. 2004. Structural and Functional Characterization of π Bulges and Other Short Intrahelical Deformations. Structure. 12(1):133–144. doi:10.1016/j.str.2003.12.001.

Cooley RB, Arp DJ, Karplus PA. 2010. Evolutionary Origin of a Secondary Structure: π-Helices as Cryptic but Widespread Insertional Variations of α-Helices That Enhance Protein Functionality. Journal of Molecular Biology. 404(2):232–246. doi:10.1016/j.jmb.2010.09.034.

Cubellis MV, Caillez F, Blundell TL, Lovell SC. 2005. Properties of polyproline II, a secondary structure element implicated in protein–protein interactions. Proteins: Structure, Function, and Bioinformatics. 58(4):880–892. doi:10.1002/prot.20327.

Frishman D, Argos P. 1995. Knowledge-based protein secondary structure assignment. Proteins: Structure, Function, and Bioinformatics. 23(4):566–579. doi:10.1002/prot.340230412.

Harris CR, Millman KJ, van der Walt SJ, Gommers R, Virtanen P, Cournapeau D, Wieser E, Taylor J, Berg S, Smith NJ, et al. 2020. Array programming with NumPy. Nature. 585(7825):357–362. doi:10.1038/s41586-020-2649-2.

Hayes T, Rao R, Akin H, Sofroniew NJ, Oktay D, Lin Z, Verkuil R, Tran VQ, Deaton J, Wiggert M, et al. 2025. Simulating 500 million years of evolution with a language model. Science. 0(0):eads0018. doi:10.1126/science.ads0018.

Hekkelman ML. 2025. PDB-REDO/libcifpp. [accessed 2025 Feb 7]. https://github.com/PDB-REDO/libcifpp.

Hekkelman ML, de Vries I, Joosten RP, Perrakis A. 2023. AlphaFill: enriching AlphaFold models with ligands and cofactors. Nat Methods. 20(2):205–213. doi:10.1038/s41592-022-01685-y.

Herrera LPT, Andreassen SN, Caroli J, Rodríguez-Espigares I, Kermani AA, Keserű GM, Kooistra AJ, Pándy-Szekeres G, Gloriam DE. 2025. GPCRdb in 2025: adding odorant receptors, data mapper, structure similarity search and models of physiological ligand complexes. Nucleic Acids Research. 53(D1):D425– D435. doi:10.1093/nar/gkae1065.

Hunter JD. 2007. Matplotlib: A 2D Graphics Environment. Computing in Science & Engineering. 9(3):90– 95. doi:10.1109/MCSE.2007.55.

Joosten RP, te Beek TAH, Krieger E, Hekkelman ML, Hooft RWW, Schneider R, Sander C, Vriend G. 2011. A series of PDB related databases for everyday needs. Nucleic Acids Research. 39(Suppl_1):D411–D419. doi:10.1093/nar/gkq1105.

Kabsch W, Sander C. 1983. Dictionary of protein secondary structure: Pattern recognition of hydrogen-bonded and geometrical features. Biopolymers. 22(12):2577–2637. doi:10.1002/bip.360221211.

Kim RS, Levy Karin E, Mirdita M, Chikhi R, Steinegger M. 2025. BFVD—a large repository of predicted viral protein structures. Nucleic Acids Research. 53(D1):D340–D347. doi:10.1093/nar/gkae1119.

King SM, Johnson WC. 1999. Assigning secondary structure from protein coordinate data. Proteins: Structure, Function, and Bioinformatics. 35(3):313–320. doi:10.1002/(SICI)1097-0134(19990515)35:3<313::AID-PROT5>3.0.CO;2-1.

Klose DP, Wallace BA, Janes RW. 2010. 2Struc: the secondary structure server. Bioinformatics. 26(20):2624–2625. doi:10.1093/bioinformatics/btq480.

Krissinel E, Henrick K. 2004. Secondary-structure matching (SSM), a new tool for fast protein structure alignment in three dimensions. Acta Cryst D. 60(12):2256–2268. doi:10.1107/S0907444904026460.

Labesse G, Colloc’h N, Pothier J, Mornon J-P. 1997. P-SEA: a new efficient assignment of secondary structure from Cα trace of proteins. Bioinformatics. 13(3):291–295. doi:10.1093/bioinformatics/13.3.291.

Law SM, Frank AT, Brooks III CL. 2014. PCASSO: A fast and efficient Cα-based method for accurately assigning protein secondary structure elements. Journal of Computational Chemistry. 35(24):1757–1761. doi:10.1002/jcc.23683.

Lazar T, Martínez-Pérez E, Quaglia F, Hatos A, Chemes LB, Iserte JA, Méndez NA, Garrone NA, Saldaño TE, Marchetti J, et al. 2021. PED in 2021: a major update of the protein ensemble database for intrinsically disordered proteins. Nucleic Acids Research. 49(D1):D404–D411. doi:10.1093/nar/gkaa1021.

Leung MR, Zeng J, Wang X, Roelofs MC, Huang W, Zenezini Chiozzi R, Hevler JF, Heck AJR, Dutcher SK, Brown A, et al. 2023. Structural specializations of the sperm tail. Cell. 186(13):2880-2896.e17. doi:10.1016/j.cell.2023.05.026.

Majumdar I, Krishna SS, Grishin NV. 2005. PALSSE: A program to delineate linear secondary structural elements from protein structures. BMC Bioinformatics. 6(1):202. doi:10.1186/1471-2105-6-202.

Mansiaux Y, Joseph AP, Gelly J-C, Brevern AG de. 2011. Assignment of PolyProline II Conformation and Analysis of Sequence – Structure Relationship. PLOS ONE. 6(3):e18401. doi:10.1371/journal.pone.0018401.

Martin J, Letellier G, Marin A, Taly J-F, de Brevern AG, Gibrat J-F. 2005. Protein secondary structure assignment revisited: a detailed analysis of different assignment methods. BMC Structural Biology. 5(1):17. doi:10.1186/1472-6807-5-17.

Meirson T, Bomze D, Markel G, Samson AO. 2020. κ-helix and the helical lock and key model: a pivotal way of looking at polyproline II. Bioinformatics. 36(12):3726–3732. doi:10.1093/bioinformatics/btaa186.

Murzin AG, Brenner SE, Hubbard T, Chothia C. 1995. SCOP: A structural classification of proteins database for the investigation of sequences and structures. Journal of Molecular Biology. 247(4):536–540. doi:10.1016/S0022-2836(05)80134-2.

Orengo C, Michie A, Jones S, Jones D, Swindells M, Thornton J. 1997. CATH – a hierarchic classification of protein domain structures. Structure. 5(8):1093–1109. doi:10.1016/S0969-2126(97)00260-8.

Pan Q, Nguyen TB, Ascher DB, Pires DEV. 2022. Systematic evaluation of computational tools to predict the effects of mutations on protein stability in the absence of experimental structures. Briefings in Bioinformatics. 23(2):bbac025. doi:10.1093/bib/bbac025.

Pavlopoulos GA, Baltoumas FA, Liu S, Selvitopi O, Camargo AP, Nayfach S, Azad A, Roux S, Call L, Ivanova NN, et al. 2023. Unraveling the functional dark matter through global metagenomics. Nature. 622(7983):594–602. doi:10.1038/s41586-023-06583-7.

Richardson JS, Richardson DC. 1989. Principles and Patterns of Protein Conformation. In: Fasman GD, editor. Prediction of Protein Structure and the Principles of Protein Conformation. Boston, MA: Springer US. p. 1–98. [accessed 2025 Jan 22]. 10.1007/978-1-4613-1571-1_1.

Taylor WR. 2001. Defining linear segments in protein structure1. Journal of Molecular Biology. 310(5):1135–1150. doi:10.1006/jmbi.2001.4817.

The pandas development team. 2020. pandas-dev/pandas: Pandas. doi:10.5281/zenodo.3509134. https://doi.org/10.5281/zenodo.3509134.

Touw WG, Baakman C, Black J, te Beek TAH, Krieger E, Joosten RP, Vriend G. 2015. A series of PDB-related databanks for everyday needs. Nucleic Acids Research. 43(D1):D364–D368. doi:10.1093/nar/gku1028.

Van der Kant R, Vriend G. 2014. Alpha-Bulges in G Protein-Coupled Receptors. International Journal of Molecular Sciences. 15(5):7841–7864. doi:10.3390/ijms15057841.

Varadi M, Anyango S, Deshpande M, Nair S, Natassia C, Yordanova G, Yuan D, Stroe O, Wood G, Laydon A, et al. 2022. AlphaFold Protein Structure Database: massively expanding the structural coverage of protein-sequence space with high-accuracy models. Nucleic Acids Research. 50(D1):D439–D444. doi:10.1093/nar/gkab1061.

Wang G, Dunbrack RL Jr. 2003. PISCES: a protein sequence culling server. Bioinformatics. 19(12):1589– 1591. doi:10.1093/bioinformatics/btg224.

Wang Y, Yang J, Hu F, Yang Y, Huang K, Zhang K. 2023. Cryo-EM reveals how the mastigoneme assembles and responds to environmental signal changes. Journal of Cell Biology. 222(12):e202301066. doi:10.1083/jcb.202301066.

Waskom ML. 2021. seaborn: statistical data visualization. Journal of Open Source Software. 6(60):3021. doi:10.21105/joss.03021.

Westbrook JD, Young JY, Shao C, Feng Z, Guranovic V, Lawson CL, Vallat B, Adams PD, Berrisford JM, Bricogne G, et al. 2022. PDBx/mmCIF Ecosystem: Foundational Semantic Tools for Structural Biology. Journal of Molecular Biology. 434(11):167599. doi:10.1016/j.jmb.2022.167599.

wwPDB consortium. 2019. Protein Data Bank: the single global archive for 3D macromolecular structure data. Nucleic Acids Research. 47(D1):D520–D528. doi:10.1093/nar/gky949.

Xia Y, Xia C-Q, Pan X, Shen H-B. 2021. GraphBind: protein structural context embedded rules learned by hierarchical graph neural networks for recognizing nucleic-acid-binding residues. Nucleic Acids Research. 49(9):e51. doi:10.1093/nar/gkab044.

