## Supplemental data 1 for "DSSP 4: FAIR annotation of protein secondary structure": supplemental_data1_dssp-extension.dic

dssp/libdssp/mmcif\_pdbx/dssp-extension.dic at trunk · PDB-REDO/dssp


Skip to content


#### Navigation Menu

### Global navigation

- Home
- Issues
- Pull requests
- Projects
- Discussions
- Codespaces
- Copilot
- Explore
- Marketplace

Loading

© 2025 GitHub, Inc.

About
Blog
Terms
Privacy
Security
Status


PDB-REDO
 /
 **dssp** 


### Navigate back to

- PDB-REDO
- dssp

- PDB-REDO

  /
- dssp

Type `/` to search

### Search code, repositories, users, issues, pull requests...

Search

Clear

Search syntax tips

 
Give feedback

### Provide feedback

We read every piece of feedback, and take your input very seriously.


Include my email address so I can be contacted

Cancel
 Submit feedback


### Saved searches

#### Use saved searches to filter your results more quickly

Name

Query

To see all available qualifiers, see our documentation.

Cancel
 Create saved search

Chat with Copilot


 


Open Copilot…

Create new...

 


Issues
 


Pull requests

 

Notifications

- Code
- Issues
  2
- Pull requests
  0
- Actions
- Projects
  0
- Wiki
- Security
- Insights
- Settings

Additional navigation options


- Code
- Issues
- Pull requests
- Actions
- Projects
- Wiki
- Security
- Insights
- Settings

You signed in with another tab or window. Reload to refresh your session.
You signed out in another tab or window. Reload to refresh your session.
You switched accounts on another tab or window. Reload to refresh your session.
 


Dismiss alert

{{ message }}

Open in github.dev
Open in a new github.dev tab
Open in codespace


#### Files

#### Files

trunk

Add file

`t`

- .github
- cmake
- doc
- libdssp

  - cmake
  - include
  - mmcif\_pdbx

    - dssp-extension.dic
  - src
  - CMakeLists.txt
- src
- test
- tools
- .dockerignore
- .gitignore
- .gitmodules
- CMakeLists.txt
- Dockerfile
- LICENSE
- README.md
- changelog

#### Breadcrumbs

1. dssp
2. /libdssp
3. /mmcif\_pdbx

/

### dssp-extension.dic

Copy path

Blame

Blame

#### Latest commit

#### History

History

1847 lines (1685 loc) · 61.6 KB

#### Breadcrumbs

1. dssp
2. /libdssp
3. /mmcif\_pdbx

/

### dssp-extension.dic

Top

#### File metadata and controls

- Code
- Blame

1847 lines (1685 loc) · 61.6 KB

Raw

1

2

3

4

5

6

7

8

9

10

11

12

13

14

15

16

17

18

19

20

21

22

23

24

25

26

27

28

29

30

31

32

33

34

35

36

37

38

39

40

41

42

43

44

45

46

47

48

49

50

51

52

53

54

55

56

57

58

59

60

61

62

63

64

65

66

67

68

69

70

71

72

73

74

75

76

77

78

79

80

81

82

83

84

85

86

87

88

89

90

91

92

93

94

95

96

97

98

99

100

101

102

103

104

105

106

107

108

109

110

111

112

113

114

115

116

117

118

119

120

121

122

123

124

125

126

127

128

129

130

131

132

133

134

135

136

137

138

139

140

141

142

143

144

145

146

147

148

149

150

151

152

153

154

155

156

157

158

159

160

161

162

163

164

165

166

167

168

169

170

171

172

173

174

175

176

177

178

179

180

181

182

183

184

185

186

187

188

189

190

191

192

193

194

195

196

197

198

199

200

201

202

203

204

205

206

207

208

209

210

211

212

213

214

215

216

217

218

219

220

221

222

223

224

225

226

227

228

229

230

231

232

233

234

235

236

237

238

239

240

241

242

243

244

245

246

247

248

249

250

251

252

253

254

255

256

257

258

259

260

261

262

263

264

265

266

267

268

269

270

271

272

273

274

275

276

277

278

279

280

281

282

283

284

285

286

287

288

289

290

291

292

293

294

295

296

297

298

299

300

301

302

303

304

305

306

307

308

309

310

311

312

313

314

315

316

317

318

319

320

321

322

323

324

325

326

327

328

329

330

331

332

333

334

335

336

337

338

339

340

341

342

343

344

345

346

347

348

349

350

351

352

353

354

355

356

357

358

359

360

361

362

363

364

365

366

367

368

369

370

371

372

373

374

375

376

377

378

379

380

381

382

383

384

385

386

387

388

389

390

391

392

393

394

395

396

397

398

399

400

401

402

403

404

405

406

407

408

409

410

411

412

413

414

415

416

417

418

419

420

421

422

423

424

425

426

427

428

429

430

431

432

433

434

435

436

437

438

439

440

441

442

443

444

445

446

447

448

449

450

451

452

453

454

455

456

457

458

459

460

461

462

463

464

465

466

467

468

469

470

471

472

473

474

475

476

477

478

479

480

481

482

483

484

485

486

487

488

489

490

491

492

493

494

495

496

497

498

499

500

501

502

503

504

505

506

507

508

509

510

511

512

513

514

515

516

517

518

519

520

521

522

523

524

525

526

527

528

529

530

531

532

533

534

535

536

537

538

539

540

541

542

543

544

545

546

547

548

549

550

551

552

553

554

555

556

557

558

559

560

561

562

563

564

565

566

567

568

569

570

571

572

573

574

575

576

577

578

579

580

581

582

583

584

585

586

587

588

589

590

591

592

593

594

595

596

597

598

599

600

601

602

603

604

605

606

607

608

609

610

611

612

613

614

615

616

617

618

619

620

621

622

623

624

625

626

627

628

629

630

631

632

633

634

635

636

637

638

639

640

641

642

643

644

645

646

647

648

649

650

651

652

653

654

655

656

657

658

659

660

661

662

663

664

665

666

667

668

669

670

671

672

673

674

675

676

677

678

679

680

681

682

683

684

685

686

687

688

689

690

691

692

693

694

695

696

697

698

699

700

701

702

703

704

705

706

707

708

709

710

711

712

713

714

715

716

717

718

719

720

721

722

723

724

725

726

727

728

729

730

731

732

733

734

735

736

737

738

739

740

741

742

743

744

745

746

747

748

749

750

751

752

753

754

755

756

757

758

759

760

761

762

763

764

765

766

767

768

769

770

771

772

773

774

775

776

777

778

779

780

781

782

783

784

785

786

787

788

789

790

791

792

793

794

795

796

797

798

799

800

801

802

803

804

805

806

807

808

809

810

811

812

813

814

815

816

817

818

819

820

821

822

823

824

825

826

827

828

829

830

831

832

833

834

835

836

837

838

839

840

841

842

843

844

845

846

847

848

849

850

851

852

853

854

855

856

857

858

859

860

861

862

863

864

865

866

867

868

869

870

871

872

873

874

875

876

877

878

879

880

881

882

883

884

885

886

887

888

889

890

891

892

893

894

895

896

897

898

899

900

901

902

903

904

905

906

907

908

909

910

911

912

913

914

915

916

917

918

919

920

921

922

923

924

925

926

927

928

929

930

931

932

933

934

935

936

937

938

939

940

941

942

943

944

945

946

947

948

949

950

951

952

953

954

955

956

957

958

959

960

961

962

963

964

965

966

967

968

969

970

971

972

973

974

975

976

977

978

979

980

981

982

983

984

985

986

987

988

989

990

991

992

993

994

995

996

997

998

999

1000

#### File: dssp-extension.dic

#### Date: 30-May-2023

##

### Items containing Secondary Structure information as written by e.g. DSSP

#

#

data\_mmcif\_pdbx-def-dssp.dic

########################################

#### Category dssp\_struct\_bridge\_pairs ##

########################################

save\_dssp\_struct\_bridge\_pairs

#

\_category.description 'dssp\_struct\_bridge\_pairs describes bridge pairs as assigned by DSSP.'

\_category.id dssp\_struct\_bridge\_pairs

\_category.mandatory\_code no

#

\_category\_key.name '\_dssp\_struct\_bridge\_pairs.id'

loop\_

\_category\_group.id 'inclusive\_group'

'struct\_group'

#

loop\_

\_category\_examples.detail

\_category\_examples.case

# - - - - - - - - - - - - - - - - - - - - - - - - - - - - - - - - - - - - - - -

;

Example taken from 1cbs, first bridge pair

;

;

\_dssp\_struct\_bridge\_pairs.id 2

\_dssp\_struct\_bridge\_pairs.label\_comp\_id ASN

\_dssp\_struct\_bridge\_pairs.label\_seq\_id 2

\_dssp\_struct\_bridge\_pairs.label\_asym\_id A

\_dssp\_struct\_bridge\_pairs.auth\_seq\_id 2

\_dssp\_struct\_bridge\_pairs.auth\_asym\_id A

\_dssp\_struct\_bridge\_pairs.pdbx\_PDB\_ins\_code ?

\_dssp\_struct\_bridge\_pairs.acceptor\_1\_label\_comp\_id PHE

\_dssp\_struct\_bridge\_pairs.acceptor\_1\_label\_seq\_id 3

\_dssp\_struct\_bridge\_pairs.acceptor\_1\_label\_asym\_id A

\_dssp\_struct\_bridge\_pairs.acceptor\_1\_auth\_seq\_id 3

\_dssp\_struct\_bridge\_pairs.acceptor\_1\_auth\_asym\_id A

\_dssp\_struct\_bridge\_pairs.acceptor\_1\_pdbx\_PDB\_ins\_code ?

\_dssp\_struct\_bridge\_pairs.acceptor\_1\_energy -0.1

\_dssp\_struct\_bridge\_pairs.acceptor\_2\_label\_comp\_id SER

\_dssp\_struct\_bridge\_pairs.acceptor\_2\_label\_seq\_id 4

\_dssp\_struct\_bridge\_pairs.acceptor\_2\_label\_asym\_id A

\_dssp\_struct\_bridge\_pairs.acceptor\_2\_auth\_seq\_id 4

\_dssp\_struct\_bridge\_pairs.acceptor\_2\_auth\_asym\_id A

\_dssp\_struct\_bridge\_pairs.acceptor\_2\_pdbx\_PDB\_ins\_code ?

\_dssp\_struct\_bridge\_pairs.acceptor\_2\_energy -0.1

\_dssp\_struct\_bridge\_pairs.donor\_1\_label\_comp\_id ASN

\_dssp\_struct\_bridge\_pairs.donor\_1\_label\_seq\_id 91

\_dssp\_struct\_bridge\_pairs.donor\_1\_label\_asym\_id A

\_dssp\_struct\_bridge\_pairs.donor\_1\_auth\_seq\_id 91

\_dssp\_struct\_bridge\_pairs.donor\_1\_auth\_asym\_id A

\_dssp\_struct\_bridge\_pairs.donor\_1\_pdbx\_PDB\_ins\_code ?

\_dssp\_struct\_bridge\_pairs.donor\_1\_energy -0.0

\_dssp\_struct\_bridge\_pairs.donor\_2\_label\_comp\_id GLU

\_dssp\_struct\_bridge\_pairs.donor\_2\_label\_seq\_id 46

\_dssp\_struct\_bridge\_pairs.donor\_2\_label\_asym\_id A

\_dssp\_struct\_bridge\_pairs.donor\_2\_auth\_seq\_id 46

\_dssp\_struct\_bridge\_pairs.donor\_2\_auth\_asym\_id A

\_dssp\_struct\_bridge\_pairs.donor\_2\_pdbx\_PDB\_ins\_code ?

\_dssp\_struct\_bridge\_pairs.donor\_2\_energy -0.0

;

save\_

#

save\_\_dssp\_struct\_bridge\_pairs.id

\_item.description

; The value of \_dssp\_struct\_bridge\_pairs.id must uniquely identify a record in the

dssp\_struct\_bridge\_pairs list.

;

\_item.name '\_dssp\_struct\_bridge\_pairs.id'

\_item.category\_id dssp\_struct\_bridge\_pairs

\_item.mandatory\_code yes

\_item\_type.code code

save\_

save\_\_dssp\_struct\_bridge\_pairs.label\_comp\_id

\_item.description

; A component of the identifier for this monomer.

This data item is a pointer to \_chem\_comp.id in the CHEM\_COMP

category.

;

\_item.name '\_dssp\_struct\_bridge\_pairs.label\_comp\_id'

\_item.category\_id dssp\_struct\_bridge\_pairs

\_item.mandatory\_code yes

\_item\_type.code ucode

save\_

save\_\_dssp\_struct\_bridge\_pairs.label\_seq\_id

\_item.description

; A component of the identifier for this monomer.

;

\_item.name '\_dssp\_struct\_bridge\_pairs.label\_seq\_id'

\_item.category\_id dssp\_struct\_bridge\_pairs

\_item.mandatory\_code yes

\_item\_type.code int

save\_

save\_\_dssp\_struct\_bridge\_pairs.label\_asym\_id

\_item.description

; A component of the identifier for this monomer

;

\_item.name '\_dssp\_struct\_bridge\_pairs.label\_asym\_id'

\_item.category\_id dssp\_struct\_bridge\_pairs

\_item.mandatory\_code yes

\_item\_type.code code

save\_

save\_\_dssp\_struct\_bridge\_pairs.auth\_seq\_id

\_item.description

; A component of the identifier for this monomer

;

\_item.name '\_dssp\_struct\_bridge\_pairs.auth\_seq\_id'

\_item.category\_id dssp\_struct\_bridge\_pairs

\_item.mandatory\_code no

\_item\_type.code int

save\_

save\_\_dssp\_struct\_bridge\_pairs.auth\_asym\_id

\_item.description

; A component of the identifier for this monomer

;

\_item.name '\_dssp\_struct\_bridge\_pairs.auth\_asym\_id'

\_item.category\_id dssp\_struct\_bridge\_pairs

\_item.mandatory\_code no

\_item\_type.code code

save\_

save\_\_dssp\_struct\_bridge\_pairs.pdbx\_PDB\_ins\_code

\_item.description

; A component of the identifier for this monomer

;

\_item.name '\_dssp\_struct\_bridge\_pairs.pdbx\_PDB\_ins\_code'

\_item.category\_id dssp\_struct\_bridge\_pairs

\_item.mandatory\_code no

\_item\_type.code code

save\_

save\_\_dssp\_struct\_bridge\_pairs.acceptor\_1\_label\_comp\_id

\_item.description

; A component of the identifier of the first residue that forms a bridge pair

with this monomer where this monomer is the donor.

;

\_item.name '\_dssp\_struct\_bridge\_pairs.acceptor\_1\_label\_comp\_id'

\_item.category\_id dssp\_struct\_bridge\_pairs

\_item.mandatory\_code no

\_item\_type.code ucode

save\_

save\_\_dssp\_struct\_bridge\_pairs.acceptor\_1\_label\_seq\_id

\_item.description

; A component of the identifier of the first residue that forms a bridge pair

with this monomer where this monomer is the donor.

;

\_item.name '\_dssp\_struct\_bridge\_pairs.acceptor\_1\_label\_seq\_id'

\_item.category\_id dssp\_struct\_bridge\_pairs

\_item.mandatory\_code no

\_item\_type.code int

save\_

save\_\_dssp\_struct\_bridge\_pairs.acceptor\_1\_label\_asym\_id

\_item.description

; A component of the identifier of the first residue that forms a bridge pair

with this monomer where this monomer is the donor.

;

\_item.name '\_dssp\_struct\_bridge\_pairs.acceptor\_1\_label\_asym\_id'

\_item.category\_id dssp\_struct\_bridge\_pairs

\_item.mandatory\_code no

\_item\_type.code code

save\_

save\_\_dssp\_struct\_bridge\_pairs.acceptor\_1\_auth\_seq\_id

\_item.description

; A component of the identifier of the first residue that forms a bridge pair

with this monomer where this monomer is the donor.

;

\_item.name '\_dssp\_struct\_bridge\_pairs.acceptor\_1\_auth\_seq\_id'

\_item.category\_id dssp\_struct\_bridge\_pairs

\_item.mandatory\_code no

\_item\_type.code int

save\_

save\_\_dssp\_struct\_bridge\_pairs.acceptor\_1\_auth\_asym\_id

\_item.description

; A component of the identifier of the first residue that forms a bridge pair

with this monomer where this monomer is the donor.

;

\_item.name '\_dssp\_struct\_bridge\_pairs.acceptor\_1\_auth\_asym\_id'

\_item.category\_id dssp\_struct\_bridge\_pairs

\_item.mandatory\_code no

\_item\_type.code code

save\_

save\_\_dssp\_struct\_bridge\_pairs.acceptor\_1\_pdbx\_PDB\_ins\_code

\_item.description

; A component of the identifier of the first residue that forms a bridge pair

with this monomer where this monomer is the donor.

;

\_item.name '\_dssp\_struct\_bridge\_pairs.acceptor\_1\_pdbx\_PDB\_ins\_code'

\_item.category\_id dssp\_struct\_bridge\_pairs

\_item.mandatory\_code no

\_item\_type.code code

save\_

save\_\_dssp\_struct\_bridge\_pairs.acceptor\_1\_energy

\_item.description

; The calculated energy for the H-bond between this residue and the first

acceptor.

;

\_item.name '\_dssp\_struct\_bridge\_pairs.acceptor\_1\_energy'

\_item.category\_id dssp\_struct\_bridge\_pairs

\_item.mandatory\_code no

\_item\_type.code float

save\_

save\_\_dssp\_struct\_bridge\_pairs.acceptor\_2\_label\_comp\_id

\_item.description

; A component of the identifier of the second residue that forms a bridge pair

with this monomer where this monomer is the donor.

;

\_item.name '\_dssp\_struct\_bridge\_pairs.acceptor\_2\_label\_comp\_id'

\_item.category\_id dssp\_struct\_bridge\_pairs

\_item.mandatory\_code no

\_item\_type.code ucode

save\_

save\_\_dssp\_struct\_bridge\_pairs.acceptor\_2\_label\_seq\_id

\_item.description

; A component of the identifier of the second residue that forms a bridge pair

with this monomer where this monomer is the donor.

;

\_item.name '\_dssp\_struct\_bridge\_pairs.acceptor\_2\_label\_seq\_id'

\_item.category\_id dssp\_struct\_bridge\_pairs

\_item.mandatory\_code no

\_item\_type.code int

save\_

save\_\_dssp\_struct\_bridge\_pairs.acceptor\_2\_label\_asym\_id

\_item.description

; A component of the identifier of the second residue that forms a bridge pair

with this monomer where this monomer is the donor.

;

\_item.name '\_dssp\_struct\_bridge\_pairs.acceptor\_2\_label\_asym\_id'

\_item.category\_id dssp\_struct\_bridge\_pairs

\_item.mandatory\_code no

\_item\_type.code code

save\_

save\_\_dssp\_struct\_bridge\_pairs.acceptor\_2\_auth\_seq\_id

\_item.description

; A component of the identifier of the second residue that forms a bridge pair

with this monomer where this monomer is the donor.

;

\_item.name '\_dssp\_struct\_bridge\_pairs.acceptor\_2\_auth\_seq\_id'

\_item.category\_id dssp\_struct\_bridge\_pairs

\_item.mandatory\_code no

\_item\_type.code int

save\_

save\_\_dssp\_struct\_bridge\_pairs.acceptor\_2\_auth\_asym\_id

\_item.description

; A component of the identifier of the second residue that forms a bridge pair

with this monomer where this monomer is the donor.

;

\_item.name '\_dssp\_struct\_bridge\_pairs.acceptor\_2\_auth\_asym\_id'

\_item.category\_id dssp\_struct\_bridge\_pairs

\_item.mandatory\_code no

\_item\_type.code code

save\_

save\_\_dssp\_struct\_bridge\_pairs.acceptor\_2\_pdbx\_PDB\_ins\_code

\_item.description

; A component of the identifier of the second residue that forms a bridge pair

with this monomer where this monomer is the donor.

;

\_item.name '\_dssp\_struct\_bridge\_pairs.acceptor\_2\_pdbx\_PDB\_ins\_code'

\_item.category\_id dssp\_struct\_bridge\_pairs

\_item.mandatory\_code no

\_item\_type.code code

save\_

save\_\_dssp\_struct\_bridge\_pairs.acceptor\_2\_energy

\_item.description

; The calculated energy for the H-bond between this residue and the second

acceptor.

;

\_item.name '\_dssp\_struct\_bridge\_pairs.acceptor\_2\_energy'

\_item.category\_id dssp\_struct\_bridge\_pairs

\_item.mandatory\_code no

\_item\_type.code float

save\_

save\_\_dssp\_struct\_bridge\_pairs.donor\_1\_label\_comp\_id

\_item.description

; A component of the identifier of the first residue that forms a bridge pair

with this monomer where this monomer is the acceptor.

;

\_item.name '\_dssp\_struct\_bridge\_pairs.donor\_1\_label\_comp\_id'

\_item.category\_id dssp\_struct\_bridge\_pairs

\_item.mandatory\_code no

\_item\_type.code ucode

save\_

save\_\_dssp\_struct\_bridge\_pairs.donor\_1\_label\_seq\_id

\_item.description

; A component of the identifier of the first residue that forms a bridge pair

with this monomer where this monomer is the acceptor.

;

\_item.name '\_dssp\_struct\_bridge\_pairs.donor\_1\_label\_seq\_id'

\_item.category\_id dssp\_struct\_bridge\_pairs

\_item.mandatory\_code no

\_item\_type.code int

save\_

save\_\_dssp\_struct\_bridge\_pairs.donor\_1\_label\_asym\_id

\_item.description

; A component of the identifier of the first residue that forms a bridge pair

with this monomer where this monomer is the acceptor.

;

\_item.name '\_dssp\_struct\_bridge\_pairs.donor\_1\_label\_asym\_id'

\_item.category\_id dssp\_struct\_bridge\_pairs

\_item.mandatory\_code no

\_item\_type.code code

save\_

save\_\_dssp\_struct\_bridge\_pairs.donor\_1\_auth\_seq\_id

\_item.description

; A component of the identifier of the first residue that forms a bridge pair

with this monomer where this monomer is the acceptor.

;

\_item.name '\_dssp\_struct\_bridge\_pairs.donor\_1\_auth\_seq\_id'

\_item.category\_id dssp\_struct\_bridge\_pairs

\_item.mandatory\_code no

\_item\_type.code int

save\_

save\_\_dssp\_struct\_bridge\_pairs.donor\_1\_auth\_asym\_id

\_item.description

; A component of the identifier of the first residue that forms a bridge pair

with this monomer where this monomer is the acceptor.

;

\_item.name '\_dssp\_struct\_bridge\_pairs.donor\_1\_auth\_asym\_id'

\_item.category\_id dssp\_struct\_bridge\_pairs

\_item.mandatory\_code no

\_item\_type.code code

save\_

save\_\_dssp\_struct\_bridge\_pairs.donor\_1\_pdbx\_PDB\_ins\_code

\_item.description

; A component of the identifier of the first residue that forms a bridge pair

with this monomer where this monomer is the acceptor.

;

\_item.name '\_dssp\_struct\_bridge\_pairs.donor\_1\_pdbx\_PDB\_ins\_code'

\_item.category\_id dssp\_struct\_bridge\_pairs

\_item.mandatory\_code no

\_item\_type.code code

save\_

save\_\_dssp\_struct\_bridge\_pairs.donor\_1\_energy

\_item.description

; The calculated energy for the H-bond between this residue and the first

donor.

;

\_item.name '\_dssp\_struct\_bridge\_pairs.donor\_1\_energy'

\_item.category\_id dssp\_struct\_bridge\_pairs

\_item.mandatory\_code no

\_item\_type.code float

save\_

save\_\_dssp\_struct\_bridge\_pairs.donor\_2\_label\_comp\_id

\_item.description

; A component of the identifier of the second residue that forms a bridge pair

with this monomer where this monomer is the acceptor.

;

\_item.name '\_dssp\_struct\_bridge\_pairs.donor\_2\_label\_comp\_id'

\_item.category\_id dssp\_struct\_bridge\_pairs

\_item.mandatory\_code no

\_item\_type.code ucode

save\_

save\_\_dssp\_struct\_bridge\_pairs.donor\_2\_label\_seq\_id

\_item.description

; A component of the identifier of the second residue that forms a bridge pair

with this monomer where this monomer is the acceptor.

;

\_item.name '\_dssp\_struct\_bridge\_pairs.donor\_2\_label\_seq\_id'

\_item.category\_id dssp\_struct\_bridge\_pairs

\_item.mandatory\_code no

\_item\_type.code int

save\_

save\_\_dssp\_struct\_bridge\_pairs.donor\_2\_label\_asym\_id

\_item.description

; A component of the identifier of the second residue that forms a bridge pair

with this monomer where this monomer is the acceptor.

;

\_item.name '\_dssp\_struct\_bridge\_pairs.donor\_2\_label\_asym\_id'

\_item.category\_id dssp\_struct\_bridge\_pairs

\_item.mandatory\_code no

\_item\_type.code code

save\_

save\_\_dssp\_struct\_bridge\_pairs.donor\_2\_auth\_seq\_id

\_item.description

; A component of the identifier of the second residue that forms a bridge pair

with this monomer where this monomer is the acceptor.

;

\_item.name '\_dssp\_struct\_bridge\_pairs.donor\_2\_auth\_seq\_id'

\_item.category\_id dssp\_struct\_bridge\_pairs

\_item.mandatory\_code no

\_item\_type.code int

save\_

save\_\_dssp\_struct\_bridge\_pairs.donor\_2\_auth\_asym\_id

\_item.description

; A component of the identifier of the second residue that forms a bridge pair

with this monomer where this monomer is the acceptor.

;

\_item.name '\_dssp\_struct\_bridge\_pairs.donor\_2\_auth\_asym\_id'

\_item.category\_id dssp\_struct\_bridge\_pairs

\_item.mandatory\_code no

\_item\_type.code code

save\_

save\_\_dssp\_struct\_bridge\_pairs.donor\_2\_pdbx\_PDB\_ins\_code

\_item.description

; A component of the identifier of the second residue that forms a bridge pair

with this monomer where this monomer is the acceptor.

;

\_item.name '\_dssp\_struct\_bridge\_pairs.donor\_2\_pdbx\_PDB\_ins\_code'

\_item.category\_id dssp\_struct\_bridge\_pairs

\_item.mandatory\_code no

\_item\_type.code code

save\_

save\_\_dssp\_struct\_bridge\_pairs.donor\_2\_energy

\_item.description

; The calculated energy for the H-bond between this residue and the second

donor.

;

\_item.name '\_dssp\_struct\_bridge\_pairs.donor\_2\_energy'

\_item.category\_id dssp\_struct\_bridge\_pairs

\_item.mandatory\_code no

\_item\_type.code float

save\_

########################################

#### Category dssp\_struct\_ladder ##

########################################

save\_dssp\_struct\_ladder

#

\_category.description 'dssp\_struct\_ladder describes ladders as assigned by DSSP.'

\_category.id dssp\_struct\_ladder

\_category.mandatory\_code no

#

\_category\_key.name '\_dssp\_struct\_ladder.id'

loop\_

\_category\_group.id 'inclusive\_group'

'struct\_group'

#

loop\_

\_category\_examples.detail

\_category\_examples.case

# - - - - - - - - - - - - - - - - - - - - - - - - - - - - - - - - - - - - - - -

;

Example taken from 1cbs, first ladder

;

;

\_dssp\_struct\_ladder.id A

\_dssp\_struct\_ladder.sheet\_id A

\_dssp\_struct\_ladder.range\_id\_1 A

\_dssp\_struct\_ladder.range\_id\_2 B

\_dssp\_struct\_ladder.type anti-parallel

\_dssp\_struct\_ladder.beg\_1\_label\_comp\_id GLY

\_dssp\_struct\_ladder.beg\_1\_label\_asym\_id A

\_dssp\_struct\_ladder.beg\_1\_label\_seq\_id 5

\_dssp\_struct\_ladder.pdbx\_beg\_1\_PDB\_ins\_code ?

\_dssp\_struct\_ladder.end\_1\_label\_comp\_id TRP

\_dssp\_struct\_ladder.end\_1\_label\_asym\_id A

\_dssp\_struct\_ladder.end\_1\_label\_seq\_id 7

\_dssp\_struct\_ladder.pdbx\_end\_1\_PDB\_ins\_code ?

\_dssp\_struct\_ladder.beg\_1\_auth\_comp\_id GLY

\_dssp\_struct\_ladder.beg\_1\_auth\_asym\_id A

\_dssp\_struct\_ladder.beg\_1\_auth\_seq\_id 5

\_dssp\_struct\_ladder.end\_1\_auth\_comp\_id TRP

\_dssp\_struct\_ladder.end\_1\_auth\_asym\_id A

\_dssp\_struct\_ladder.end\_1\_auth\_seq\_id 7

\_dssp\_struct\_ladder.beg\_2\_label\_comp\_id ILE

\_dssp\_struct\_ladder.beg\_2\_label\_asym\_id A

\_dssp\_struct\_ladder.beg\_2\_label\_seq\_id 43

\_dssp\_struct\_ladder.pdbx\_beg\_2\_PDB\_ins\_code ?

\_dssp\_struct\_ladder.end\_2\_label\_comp\_id VAL

\_dssp\_struct\_ladder.end\_2\_label\_asym\_id A

\_dssp\_struct\_ladder.end\_2\_label\_seq\_id 41

\_dssp\_struct\_ladder.pdbx\_end\_2\_PDB\_ins\_code ?

\_dssp\_struct\_ladder.beg\_2\_auth\_comp\_id ILE

\_dssp\_struct\_ladder.beg\_2\_auth\_asym\_id A

\_dssp\_struct\_ladder.beg\_2\_auth\_seq\_id 43

\_dssp\_struct\_ladder.end\_2\_auth\_comp\_id VAL

\_dssp\_struct\_ladder.end\_2\_auth\_asym\_id A

\_dssp\_struct\_ladder.end\_2\_auth\_seq\_id 41

;

save\_

save\_\_dssp\_struct\_ladder.id

\_item.description

; The value of \_dssp\_struct\_ladder.id must uniquely identify a record in the

dssp\_struct\_ladder list.

;

\_item.name '\_dssp\_struct\_ladder.id'

\_item.category\_id dssp\_struct\_ladder

\_item.mandatory\_code yes

\_item\_type.code code

save\_

save\_\_dssp\_struct\_ladder.sheet\_id

\_item.description

; This data item is a pointer to \_struct\_sheet.id in the

STRUCT\_SHEET category.

;

\_item.name '\_dssp\_struct\_ladder.sheet\_id'

\_item.category\_id dssp\_struct\_ladder

\_item.mandatory\_code yes

\_item\_type.code code

save\_

save\_\_dssp\_struct\_ladder.range\_id\_1

\_item.description

; This data item is a pointer to \_struct\_sheet\_range.id in

the STRUCT\_SHEET\_RANGE category.

;

\_item.name '\_dssp\_struct\_ladder.range\_id\_1'

\_item.category\_id dssp\_struct\_ladder

\_item.mandatory\_code yes

\_item\_type.code code

save\_

save\_\_dssp\_struct\_ladder.range\_id\_2

\_item.description

; This data item is a pointer to \_struct\_sheet\_range.id in

the STRUCT\_SHEET\_RANGE category.

;

\_item.name '\_dssp\_struct\_ladder.range\_id\_2'

\_item.category\_id dssp\_struct\_ladder

\_item.mandatory\_code yes

\_item\_type.code code

save\_

save\_\_dssp\_struct\_ladder.type

\_item.description

; The type of the ladder, be it parallel or anti-parallel

;

\_item.name '\_dssp\_struct\_ladder.type'

\_item.category\_id dssp\_struct\_ladder

\_item.mandatory\_code yes

\_item\_type.code code

loop\_

\_item\_enumeration.value

\_item\_enumeration.detail

parallel 'The strands forming a ladder are oriented parallel'

anti-parallel 'The strands forming a ladder are oriented anti-parallel'

save\_

save\_\_dssp\_struct\_ladder.beg\_1\_label\_comp\_id

\_item.description

; A component of the identifier for the residue at which the

first ladder segment begins.

;

\_item.name '\_dssp\_struct\_ladder.beg\_1\_label\_comp\_id'

\_item.category\_id dssp\_struct\_ladder

\_item.mandatory\_code yes

\_item\_type.code ucode

save\_

save\_\_dssp\_struct\_ladder.beg\_1\_label\_asym\_id

\_item.description

; A component of the identifier for the residue at which the

first ladder segment begins.

;

\_item.name '\_dssp\_struct\_ladder.beg\_1\_label\_asym\_id'

\_item.category\_id dssp\_struct\_ladder

\_item.mandatory\_code yes

\_item\_type.code code

save\_

save\_\_dssp\_struct\_ladder.beg\_1\_label\_seq\_id

\_item.description

; A component of the identifier for the residue at which the

first ladder segment begins.

;

\_item.name '\_dssp\_struct\_ladder.beg\_1\_label\_seq\_id'

\_item.category\_id dssp\_struct\_ladder

\_item.mandatory\_code yes

\_item\_type.code int

save\_

save\_\_dssp\_struct\_ladder.pdbx\_beg\_1\_PDB\_ins\_code

\_item.description

; A component of the identifier for the residue at which the

first ladder segment begins.

;

\_item.name '\_dssp\_struct\_ladder.pdbx\_beg\_1\_PDB\_ins\_code'

\_item.category\_id dssp\_struct\_ladder

\_item.mandatory\_code no

\_item\_type.code code

save\_

save\_\_dssp\_struct\_ladder.end\_1\_label\_comp\_id

\_item.description

; A component of the identifier for the residue at which the

first ladder segment ends.

;

\_item.name '\_dssp\_struct\_ladder.end\_1\_label\_comp\_id'

\_item.category\_id dssp\_struct\_ladder

\_item.mandatory\_code yes

\_item\_type.code ucode

save\_

save\_\_dssp\_struct\_ladder.end\_1\_label\_asym\_id

\_item.description

; A component of the identifier for the residue at which the

first ladder segment ends.

;

\_item.name '\_dssp\_struct\_ladder.end\_1\_label\_asym\_id'

\_item.category\_id dssp\_struct\_ladder

\_item.mandatory\_code yes

\_item\_type.code code

save\_

save\_\_dssp\_struct\_ladder.end\_1\_label\_seq\_id

\_item.description

; A component of the identifier for the residue at which the

first ladder segment ends.

;

\_item.name '\_dssp\_struct\_ladder.end\_1\_label\_seq\_id'

\_item.category\_id dssp\_struct\_ladder

\_item.mandatory\_code yes

\_item\_type.code int

save\_

save\_\_dssp\_struct\_ladder.pdbx\_end\_1\_PDB\_ins\_code

\_item.description

; A component of the identifier for the residue at which the

first ladder segment ends.

;

\_item.name '\_dssp\_struct\_ladder.pdbx\_end\_1\_PDB\_ins\_code'

\_item.category\_id dssp\_struct\_ladder

\_item.mandatory\_code no

\_item\_type.code code

save\_

save\_\_dssp\_struct\_ladder.beg\_1\_auth\_comp\_id

\_item.description

; A component of the identifier for the residue at which the

first ladder segment begins.

;

\_item.name '\_dssp\_struct\_ladder.beg\_1\_auth\_comp\_id'

\_item.category\_id dssp\_struct\_ladder

\_item.mandatory\_code no

\_item\_type.code ucode

save\_

save\_\_dssp\_struct\_ladder.beg\_1\_auth\_asym\_id

\_item.description

; A component of the identifier for the residue at which the

first ladder segment begins.

;

\_item.name '\_dssp\_struct\_ladder.beg\_1\_auth\_asym\_id'

\_item.category\_id dssp\_struct\_ladder

\_item.mandatory\_code no

\_item\_type.code code

save\_

save\_\_dssp\_struct\_ladder.beg\_1\_auth\_seq\_id

\_item.description

; A component of the identifier for the residue at which the

first ladder segment begins.

;

\_item.name '\_dssp\_struct\_ladder.beg\_1\_auth\_seq\_id'

\_item.category\_id dssp\_struct\_ladder

\_item.mandatory\_code no

\_item\_type.code int

save\_

save\_\_dssp\_struct\_ladder.end\_1\_auth\_comp\_id

\_item.description

; A component of the identifier for the residue at which the

first ladder segment ends.

;

\_item.name '\_dssp\_struct\_ladder.end\_1\_auth\_comp\_id'

\_item.category\_id dssp\_struct\_ladder

\_item.mandatory\_code no

\_item\_type.code ucode

save\_

save\_\_dssp\_struct\_ladder.end\_1\_auth\_asym\_id

\_item.description

; A component of the identifier for the residue at which the

first ladder segment ends.

;

\_item.name '\_dssp\_struct\_ladder.end\_1\_auth\_asym\_id'

\_item.category\_id dssp\_struct\_ladder

\_item.mandatory\_code no

\_item\_type.code code

save\_

save\_\_dssp\_struct\_ladder.end\_1\_auth\_seq\_id

\_item.description

; A component of the identifier for the residue at which the

first ladder segment ends.

;

\_item.name '\_dssp\_struct\_ladder.end\_1\_auth\_seq\_id'

\_item.category\_id dssp\_struct\_ladder

\_item.mandatory\_code no

\_item\_type.code int

save\_

save\_\_dssp\_struct\_ladder.beg\_2\_label\_comp\_id

\_item.description

; A component of the identifier for the residue at which the

second ladder segment begins.

;

\_item.name '\_dssp\_struct\_ladder.beg\_2\_label\_comp\_id'

\_item.category\_id dssp\_struct\_ladder

\_item.mandatory\_code yes

\_item\_type.code ucode

save\_

save\_\_dssp\_struct\_ladder.beg\_2\_label\_asym\_id

\_item.description

; A component of the identifier for the residue at which the

second ladder segment begins.

;

\_item.name '\_dssp\_struct\_ladder.beg\_2\_label\_asym\_id'

\_item.category\_id dssp\_struct\_ladder

\_item.mandatory\_code yes

\_item\_type.code code

save\_

save\_\_dssp\_struct\_ladder.beg\_2\_label\_seq\_id

\_item.description

; A component of the identifier for the residue at which the

second ladder segment begins.

;

\_item.name '\_dssp\_struct\_ladder.beg\_2\_label\_seq\_id'

\_item.category\_id dssp\_struct\_ladder

\_item.mandatory\_code yes

\_item\_type.code int

save\_

save\_\_dssp\_struct\_ladder.pdbx\_beg\_2\_PDB\_ins\_code

\_item.description

; A component of the identifier for the residue at which the

second ladder segment begins.

;

\_item.name '\_dssp\_struct\_ladder.pdbx\_beg\_2\_PDB\_ins\_code'

\_item.category\_id dssp\_struct\_ladder

\_item.mandatory\_code no

\_item\_type.code code

save\_

save\_\_dssp\_struct\_ladder.end\_2\_label\_comp\_id

\_item.description

; A component of the identifier for the residue at which the

second ladder segment ends.

;

\_item.name '\_dssp\_struct\_ladder.end\_2\_label\_comp\_id'

\_item.category\_id dssp\_struct\_ladder

\_item.mandatory\_code yes

\_item\_type.code ucode

save\_

save\_\_dssp\_struct\_ladder.end\_2\_label\_asym\_id

\_item.description

; A component of the identifier for the residue at which the

second ladder segment ends.

;

\_item.name '\_dssp\_struct\_ladder.end\_2\_label\_asym\_id'

\_item.category\_id dssp\_struct\_ladder

\_item.mandatory\_code yes

\_item\_type.code code

save\_

save\_\_dssp\_struct\_ladder.end\_2\_label\_seq\_id

\_item.description

; A component of the identifier for the residue at which the

second ladder segment ends.

;

\_item.name '\_dssp\_struct\_ladder.end\_2\_label\_seq\_id'

\_item.category\_id dssp\_struct\_ladder

\_item.mandatory\_code yes

\_item\_type.code int

save\_

save\_\_dssp\_struct\_ladder.pdbx\_end\_2\_PDB\_ins\_code

\_item.description

; A component of the identifier for the residue at which the

second ladder segment ends.

;

\_item.name '\_dssp\_struct\_ladder.pdbx\_end\_2\_PDB\_ins\_code'

\_item.category\_id dssp\_struct\_ladder

\_item.mandatory\_code no

\_item\_type.code code

save\_

save\_\_dssp\_struct\_ladder.beg\_2\_auth\_comp\_id

\_item.description

; A component of the identifier for the residue at which the

second ladder segment begins.

;

\_item.name '\_dssp\_struct\_ladder.beg\_2\_auth\_comp\_id'

\_item.category\_id dssp\_struct\_ladder

\_item.mandatory\_code no

\_item\_type.code ucode

save\_

save\_\_dssp\_struct\_ladder.beg\_2\_auth\_asym\_id

\_item.description

; A component of the identifier for the residue at which the

second ladder segment begins.

;

\_item.name '\_dssp\_struct\_ladder.beg\_2\_auth\_asym\_id'

\_item.category\_id dssp\_struct\_ladder

\_item.mandatory\_code no

\_item\_type.code code

save\_

save\_\_dssp\_struct\_ladder.beg\_2\_auth\_seq\_id

\_item.description

; A component of the identifier for the residue at which the

second ladder segment begins.

;

\_item.name '\_dssp\_struct\_ladder.beg\_2\_auth\_seq\_id'

\_item.category\_id dssp\_struct\_ladder

\_item.mandatory\_code no

\_item\_type.code int

save\_

save\_\_dssp\_struct\_ladder.end\_2\_auth\_comp\_id

\_item.description

; A component of the identifier for the residue at which the

second ladder segment ends.

;

\_item.name '\_dssp\_struct\_ladder.end\_2\_auth\_comp\_id'

\_item.category\_id dssp\_struct\_ladder

\_item.mandatory\_code no

\_item\_type.code ucode

save\_

save\_\_dssp\_struct\_ladder.end\_2\_auth\_asym\_id

\_item.description

; A component of the identifier for the residue at which the

second ladder segment ends.

;

\_item.name '\_dssp\_struct\_ladder.end\_2\_auth\_asym\_id'

\_item.category\_id dssp\_struct\_ladder

\_item.mandatory\_code no

\_item\_type.code code

save\_

save\_\_dssp\_struct\_ladder.end\_2\_auth\_seq\_id

\_item.description

; A component of the identifier for the residue at which the

second ladder segment ends.

;

\_item.name '\_dssp\_struct\_ladder.end\_2\_auth\_seq\_id'

\_item.category\_id dssp\_struct\_ladder

\_item.mandatory\_code no

\_item\_type.code int

save\_

########################################

#### Category dssp\_statistics ##

########################################

save\_dssp\_statistics

#

\_category.description 'dssp\_statistics contains secondary structure statistics as calculated by DSSP.'

\_category.id dssp\_statistics

\_category.mandatory\_code no

#

\_category\_key.name '\_dssp\_statistics.entry\_id'

loop\_

\_category\_group.id 'inclusive\_group'

'struct\_group'

#

loop\_

\_category\_examples.detail

\_category\_examples.case

# - - - - - - - - - - - - - - - - - - - - - - - - - - - - - - - - - - - - - - -

;

Example taken from 1cbs

;

;

\_dssp\_statistics.entry\_id 1CBS

\_dssp\_statistics.nr\_of\_residues 137

\_dssp\_statistics.nr\_of\_chains 1

\_dssp\_statistics.nr\_of\_ss\_bridges\_total 0

\_dssp\_statistics.nr\_of\_ss\_bridges\_intra\_chain 0

\_dssp\_statistics.nr\_of\_ss\_bridges\_inter\_chain 0

\_dssp\_statistics.accessible\_surface\_of\_protein 7905.01

;

save\_

save\_\_dssp\_statistics.entry\_id

\_item.description

; The entry ID for which these statistics are calculated.

;

\_item.name '\_dssp\_statistics.entry\_id'

\_item.category\_id dssp\_statistics

\_item.mandatory\_code yes

\_item\_type.code code

#

\_item\_linked.child\_name "\_dssp\_statistics.entry\_id"

\_item\_linked.parent\_name "\_entry.id"

#

save\_

save\_\_dssp\_statistics.nr\_of\_residues

\_item.description

; The total number of residues to which secondary structure information was assigned.

;

\_item.name '\_dssp\_statistics.nr\_of\_residues'

\_item.category\_id dssp\_statistics

\_item.mandatory\_code yes

\_item\_type.code int

save\_

save\_\_dssp\_statistics.nr\_of\_chains

\_item.description

; The number of chains used

;

\_item.name '\_dssp\_statistics.nr\_of\_chains'

\_item.category\_id dssp\_statistics

\_item.mandatory\_code yes

\_item\_type.code int

save\_

save\_\_dssp\_statistics.nr\_of\_ss\_bridges\_total

\_item.description

; The number of sulfur-sulfur bridges assigned.

;

\_item.name '\_dssp\_statistics.nr\_of\_ss\_bridges\_total'

\_item.category\_id dssp\_statistics

\_item.mandatory\_code yes

\_item\_type.code int

save\_

save\_\_dssp\_statistics.nr\_of\_ss\_bridges\_intra\_chain

\_item.description

; The number of intra-chain sulfur-sulfur bridges assigned.

;

\_item.name '\_dssp\_statistics.nr\_of\_ss\_bridges\_intra\_chain'

\_item.category\_id dssp\_statistics

\_item.mandatory\_code yes

\_item\_type.code int

save\_

save\_\_dssp\_statistics.nr\_of\_ss\_bridges\_inter\_chain

\_item.description

; The number of inter-chain sulfur-sulfur bridges assigned.

;

\_item.name '\_dssp\_statistics.nr\_of\_ss\_bridges\_inter\_chain'

\_item.category\_id dssp\_statistics

\_item.mandatory\_code yes

\_item\_type.code int

save\_

save\_\_dssp\_statistics.accessible\_surface\_of\_protein

\_item.description

; The total accessible surface of the protein in square Angstroms.

;

\_item.name '\_dssp\_statistics.accessible\_surface\_of\_protein'

\_item.category\_id dssp\_statistics

\_item.mandatory\_code yes

\_item\_type.code float

save\_

########################################

#### Category dssp\_statistics\_hbond ##

########################################

save\_dssp\_statistics\_hbond

#

\_category.description 'dssp\_statistics\_hbond contains statistics for the collection of h-bond variants calculated by DSSP.'

\_category.id dssp\_statistics\_hbond

\_category.mandatory\_code no

#

View remainder of file in raw view

#### Footer

© 2025 GitHub, Inc.

##### Footer navigation

- Terms
- Privacy
- Security
- Status
- Docs
- Contact
- Manage cookies
- Do not share my personal information


You can’t perform that action at this time.
